# Subtle, but Perceptible, Sexual Dichromatism and Disassortative Mating Based on Plumage Reflectance in Black Terns (*Chlidonias niger*)

**DOI:** 10.1101/2024.01.11.575209

**Authors:** Daniel T. Baldassarre, Kristina M. Davis, David A. Shealer

## Abstract

In birds, sexual selection on plumage often leads to sexual dichromatism and male ornamentation. However, even in drab species with no obvious sexual dichromatism, both sexes may still use plumage for mate choice. A previous study found weak sexual size dimorphism in North American Black Terns (*Chlidonias niger surinamensis*), but no assortative mating based on morphology. However, the relevance of plumage variation to mate choice is yet untested. Here, using reflectance spectrometry and avian visual modeling revealed that Black Tern males and females exhibited a subtle but significant difference in brightness (males darker). Importantly, the achromatic contrast between the sexes should be perceptible during mate choice. Moreover, there was evidence of disassortative mating for plumage color, but not brightness: more black and saturated birds paired with more gray and unsaturated birds. There was no relationship between plumage color or brightness and body mass. This finding suggests that visual signals unrelated to body condition may be relevant to Black Tern mate choice. The pattern of disassortative mating was unexpected, and potential benefits of choosing a mate dissimilar from oneself are discussed. This study highlights the importance of considering the avian visual system when studying plumage variation elusive to human observers.

## BLACK TERN PLUMAGE REFLECTANCE

Sexual selection often results in sexual dimorphism, particularly in polygynous species where males function primarily as sperm donors and females choose which male to mate with based on some outward phenotypic expression (Darwin 1871). In such cases, males tend to be larger, more brightly colored, or may produce some elaborate sexual ornament that serves no practical function other than to attract a female mate. Secondary sexual characteristics are common in many females, however, suggesting that mate choice is not entirely female driven, and that males also exhibit preference under certain circumstances (Amudsen 2000; Tobias *et al*. 2012). In Crested Auklets (*Aethia cristatella*), for example, mutual sexual selection may account for female ornamentation, as males and females each prefer mates with more elaborate ornaments (Jones and Hunter 1993). A similar pattern is exhibited by Black Swans (*Cygnus atratus*), where males and females mate assortatively based on the number of ornamental curled feathers (Kraaijeveld *et al*. 2004). In other species, however, female ornamentation cannot be explained by mutual mate choice, such as in the Turquoise-browed Motmot (*Eumomota superciliosa*), which does not mate assortatively based on tail ornaments, body size, or phenotypic condition (Murphy 2008).

Theoretical and empirical research on mate choice in birds has focused largely on species with obvious sexual dimorphism (Rosenthal 2017). Monomorphic species have received considerably less attention (but see Jones and Hunter 1993; Mennill *et al*. 2003; Kraaijeveld *et al*. 2004; Safran and McGraw 200; Murphy 2008; van Rooij and Griffith 2012), although they account for over half the world’s bird species (Griffiths *et al*. 1998). Moreover, the simple fact that the sexes in monomorphic species – by definition – do not exhibit any obvious secondary sexual character differences, suggests that the relevant mate choice criteria may be elusive to human observers. Most gulls and terns (Laridae) lack obvious secondary sexual characteristics, are monomorphic with respect to plumage, but may exhibit some degree of sexual size dimorphism, with males generally larger than females (e.g., Chardine and Morris 1989; Bosch 1996; Fletcher and Hamer 2003). Although size-assortative mating has been reported in some larids (Coulter 1986; Chardine and Morris 1989; Catry *et al*. 1999; Helfenstein *et al*. 2004; Ludwig and Becker 2008; but see Phillips *et al*. 2002; Nisbet *et al*. 2007), the underlying mechanisms driving the process are not well understood. For example, patterns of size-assortative mating may be confounded with age (Nisbet *et al*. 2007), particularly if males and females recruit to the breeding population at similar ages and form long-term pair bonds (Reid 1988) and if structural sizes of adults change predictably as they age (Bridge and Nisbet 2004).

A previous study (Shealer 2014) found small but significant sexual size dimorphism between male and female Black Terns (*Chlidonias niger*), with males averaging 1-5% larger in the morphometric characteristics measured. However, the same study found only weak evidence for size-assortative mating, suggesting that if Black Terns engage in mutual mate choice, the criteria may be more strongly related to behavioral traits displayed during courtship, or more cryptic traits, such as feather condition or reflectance (Shealer 2014). Previous research on the North American subspecies of Black Tern (*C. niger surinamensis*) indicates that they are both socially and genetically monogamous (Shealer *et al*. 2014), with males contributing substantially to parental effort, often incubating entirely by themselves overnight while females depart the colony for a communal roost (Custer and Custer 1996; Van der Winden 2005). Between-year mate retention is low (∼20%), however (Shealer *et al*. 2014), suggesting that most individuals must undergo the mate choice process every few years. If so, there should be strong selective pressure on both sexes to be able to discern variation in phenotypic and/or genetic quality in a prospective mate. Thus, Black Terns provide an excellent opportunity to explore the potential role of subtle plumage variation on patterns of mutual mate choice.

Here, we used reflectance spectrometry to test for sexual dimorphism in feather hue, saturation, and brightness between adult male and female Black Terns. We also used these feather characteristics to test for assortative mating in a smaller sample of mated pairs. Reflectance spectrometry has been widely used to examine patterns of mate choice, primarily in species in which males are brightly colored and exhibit chromatic variation (but see Mennill *et al*. 2003; Enbody *et al*. 2017; Taff *et al*. 2019; Feldmann *et al*. 2021). We are unaware of any published study that has used this technique to examine assortative mating in species with mainly achromatic feathers (i.e., white, black). We also explore the question of whether achromatic variation could be a sexually selected trait and if so, what the information content of the signal might be.

## METHODS

### Study Area and Sample Collection

We collected blood and breast feather samples from adult Black Terns trapped on their nests during the incubation period (under US BBL permit #22827 and appropriate letters of authorization). At the time of capture, we weighed birds to the nearest 0.5 g using a spring scale. We collected most samples (male *n* =38, female *n* = 27, including feathers from 12 mated pairs) at various colony sites in Wisconsin in 2010, except for two birds (mated pair) that we captured in 2012 at a breeding site in north central Iowa. We stored blood samples in lysis buffer for later sex determination by molecular markers (Griffiths *et al*. 1998; Fridolfsson and Ellegren 1999). We stored feather samples in zippered plastic bags, labeled by band number.

### Reflectance Spectrometry

We mounted ten feathers in an overlapping pattern on black construction paper (Strathmore Artagain Coal Black) and measured ultraviolet (UV) and visible light reflectance from 300–700 nm using an Ocean Optics Jaz reflectance spectrometer and a PX2 lamp. We inserted the probe at a 90° angle in a block that held the probe tip 5 mm from the feather surface and excluded ambient light. Between each sample, we calibrated the spectrometer using a WS-1 white standard. We measured reflectance at three different regions of the feather sample and averaged the reflectance curves before analysis.

### Avian Visual Modeling

We analyzed reflectance curves using the program *pavo 2* (Maia *et al*. 2019) in R 3.6.0 (R Core Team 2019). We obtained quantum catch values for each of the four avian single cones using the Wedge-tailed Shearwater (*Puffinus pacificus*) visual model (plus double-cone sensitivity for luminance) and idealized illuminant D65 (Hart 2004). We chose the Wedge-tailed Shearwater because it represents the closest relative to the Black Tern for which we have photoreceptor sensitivity data. Further, both species exhibit a VS-sensitive visual system (i.e., the shortest wavelength cone is stimulated primarily in the visible, not ultraviolet part of the spectrum, Ödeen *et al*. 2010). We then used the quantum catch values to plot each color as a point in tetrahedral color space where the vertices represented stimulation of the four single cones. From the color space, we extracted the angle of the color vector relative to the x-y (theta) and z (phi) plane, which together describe hue. To quantify saturation, we measured the distance between the point and the achromatic center relative to the maximum possible distance. Together these measurements described hue, saturation, and luminance of the feathers accounting for UV reflectance as perceived by the avian visual system.

We further explored whether any chromatic or achromatic differences between males and females were sufficient to be distinguished by the signal receiver. To do this, we input the quantum catch values into the noise-weighted color distance calculator in *pavo 2* using the following settings: noise: neural; photoreceptor densities: u = 1, s = 2, m = 2, l = 4; Weber fractions: 0.1. The results of this calculation are indices of either chromatic (dS) or achromatic (dL) contrast, which we compared to a value of 1, or the perceptual threshold indicating signals that can be distinguished by the avian visual system (Vorobyev and Osorio 1998).

### Statistical Analyses

Preliminary analyses revealed correlations among several of the colorspace variables, so we conducted a principal component analysis (PCA) to reduce them to a smaller number of orthogonal variables. We then compared the average male and female principal component scores using a two-sample t-test. To determine whether the noise-weighted contrast values between the sexes were larger than the avian perceptual threshold, we tested whether the average pairwise value of all inter-sex comparisons was significantly greater than 1 using a one-sample t-test. We also calculated the 95% confidence interval (CI) for the contrast values and checked for overlap with a value of 1. To explore the possibility that individuals mate assortatively based on color or brightness, we compared male and female principal component scores within mated pairs using a Pearson correlation. Finally, to evaluate the potential value of plumage color as a signal, we used a Pearson correlation to compare individual mass to the plumage variables produced by PCA. All analyses were conducted in R 3.6.0. (R Core Team 2019).

## RESULTS

Male and female breast feathers produced qualitatively similar reflectance curves with substantial overlap, although the average male curve was slightly lower (i.e., the feathers reflected less light) than the average female curve (Fig. 1).

**Figure 1:**
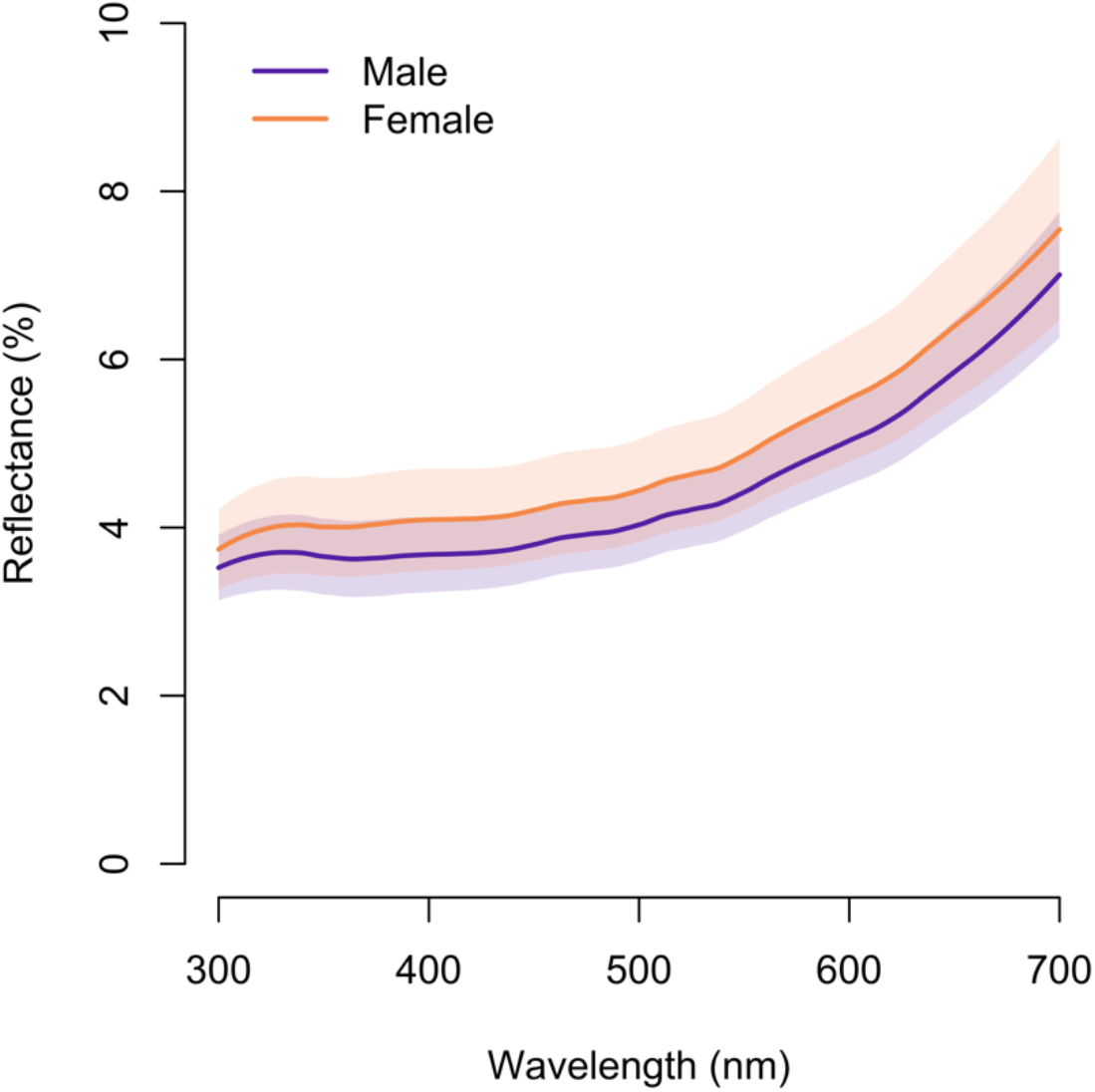
The average male and female breast feather reluctance curves (shaded area = 1 SD).

PCA resulted in two meaningful variables that we used in subsequent analyses (Table 1). PC1 (variance explained = 0.72) was loaded primarily by theta, phi, and saturation; and thus served as a chromatic measure of hue and saturation. PC2 (variance explained = 0.24) was loaded primarily by luminance, and thus served as an achromatic measure of brightness. Visualization of the PCA indicated that males and females had extensive overlap along both axes, although there was some degree of separation along PC2 (Fig. 2).

**Table 1:** Results of a principal component analysis of colorspace variables. Significant factor loadings (>0.5) are highlighted in bold.

|  | PC1 | PC2 |
| --- | --- | --- |
| Eigenvalue | 2.87 | 0.96 |
| Variance explained | 0.72 | 0.24 |
| Hue (theta) | **-0.56** | 0.21 |
| Hue (phi) | **0.59** | -1.10 |
| Saturation | **0.56** | 0.01 |
| Luminance | -0.18 | **-0.97** |

**Figure 2:**
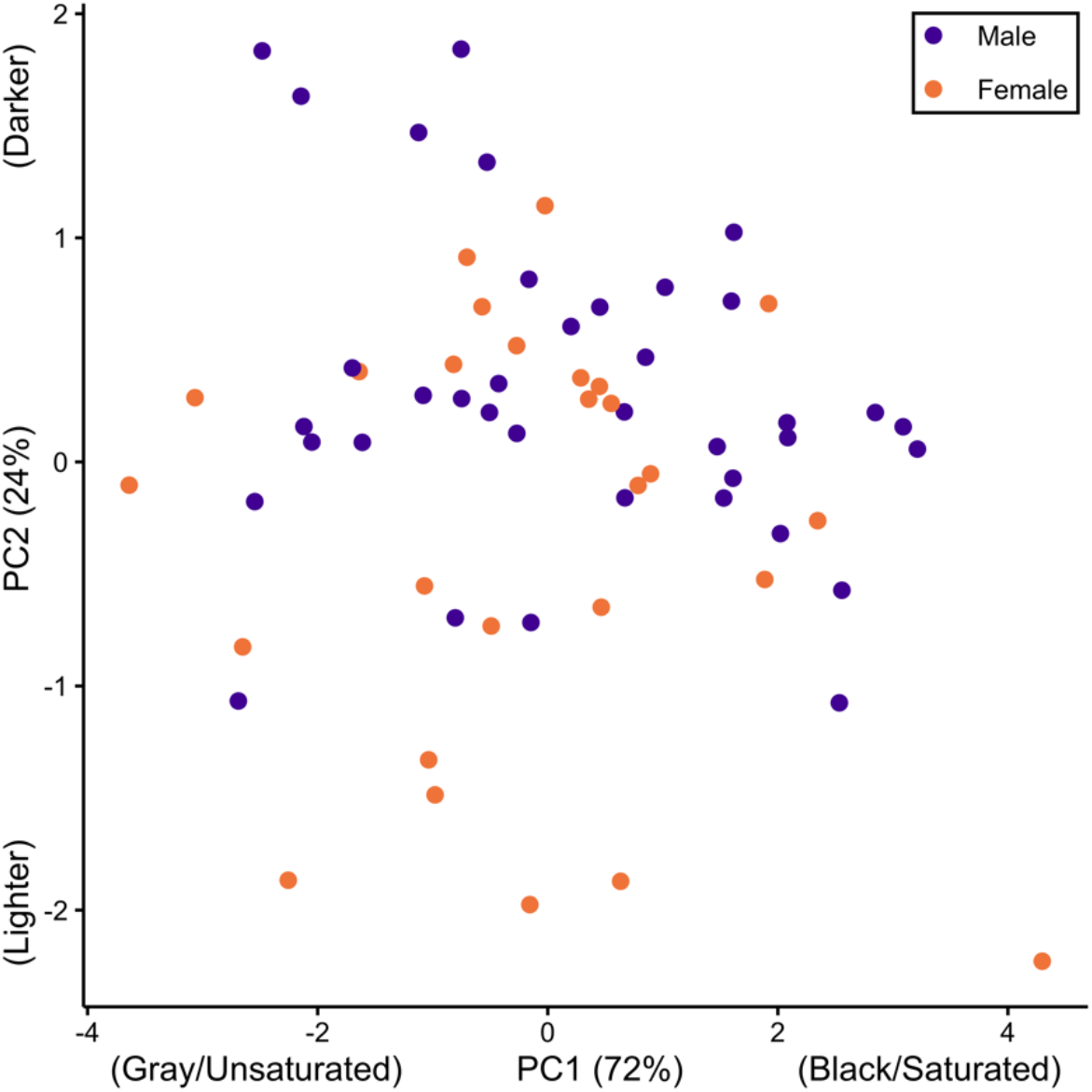
A comparison of male and female breast feather color using the first two components of a principal component analysis of colorspace variable. Numbers in parentheses along the axes indicate how much variation was explained by each component.

Supporting the qualitative inspection of the PCA plot, there was no significant difference between males and females in the average chromatic PC1 score (t_56.65_ = -0.89, *P* = 0.38), but the average male achromatic PC2 score was significantly higher (indicating darker feathers) than that of females (t_45.26_ = -2.76, *P* = 0.01; Fig. 3). Relatedly, the average chromatic contrast between males and females was not significantly higher than the perceptual threshold (<u>x</u> = 0.22, 95% CI [0.21–0.23], t_1,025_ = -156.22, *P* > 0.95), while the average achromatic contrast between males and females was significantly higher than the perceptual threshold (<u>x</u> = 1.51, 95% CI [1.44–1.58], t_1,025_ = 14.14, *P* < 0.001; Fig. 4).

**Figure 3:**
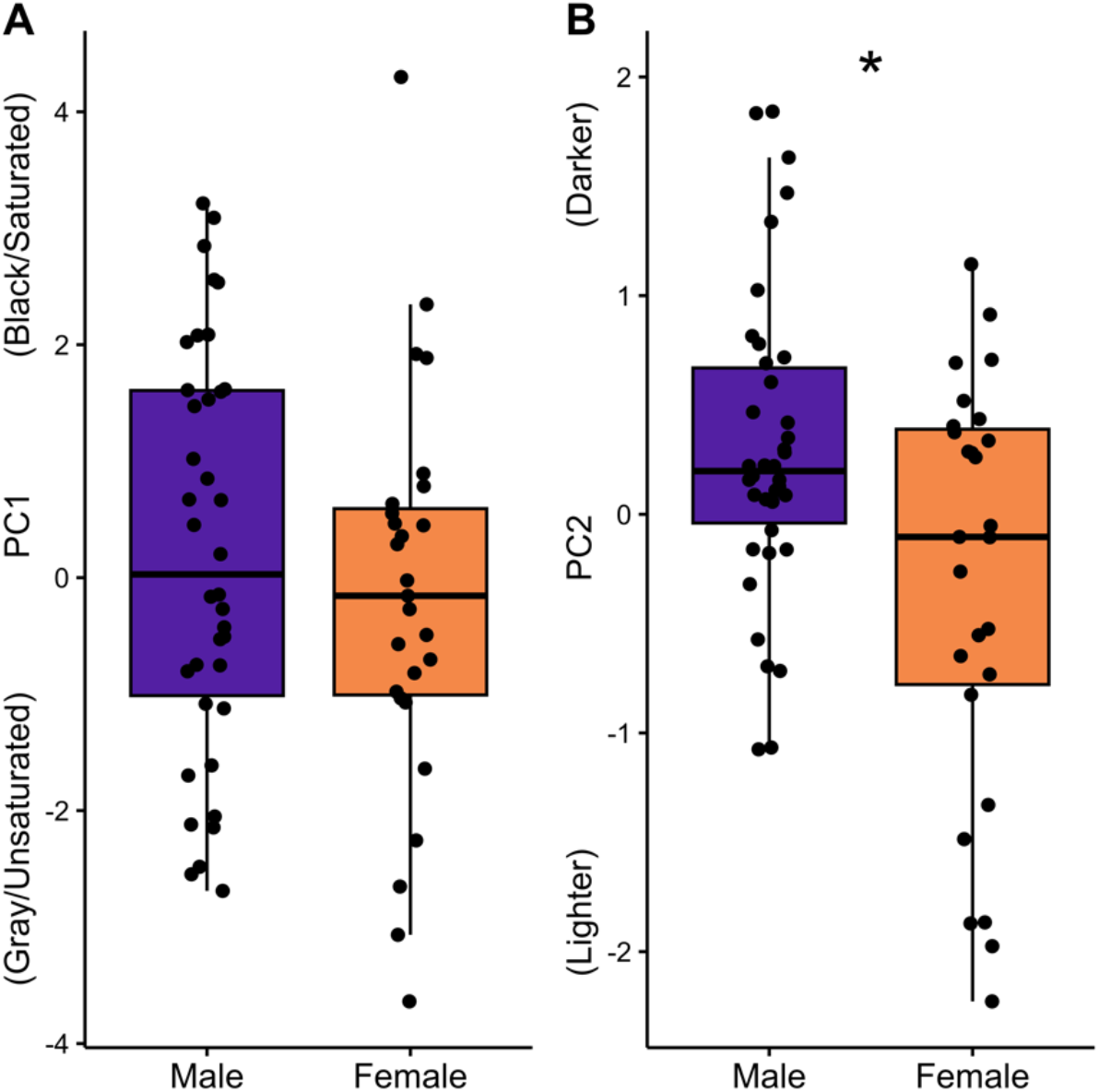
Male and female breast feathers compared using the chromatic PC1 score composed primarily of hue and saturation (A) and the achromatic PC2 score composed primarily of luminance (B). An asterisk indicates a significant difference between the sexes.

**Figure 4:**
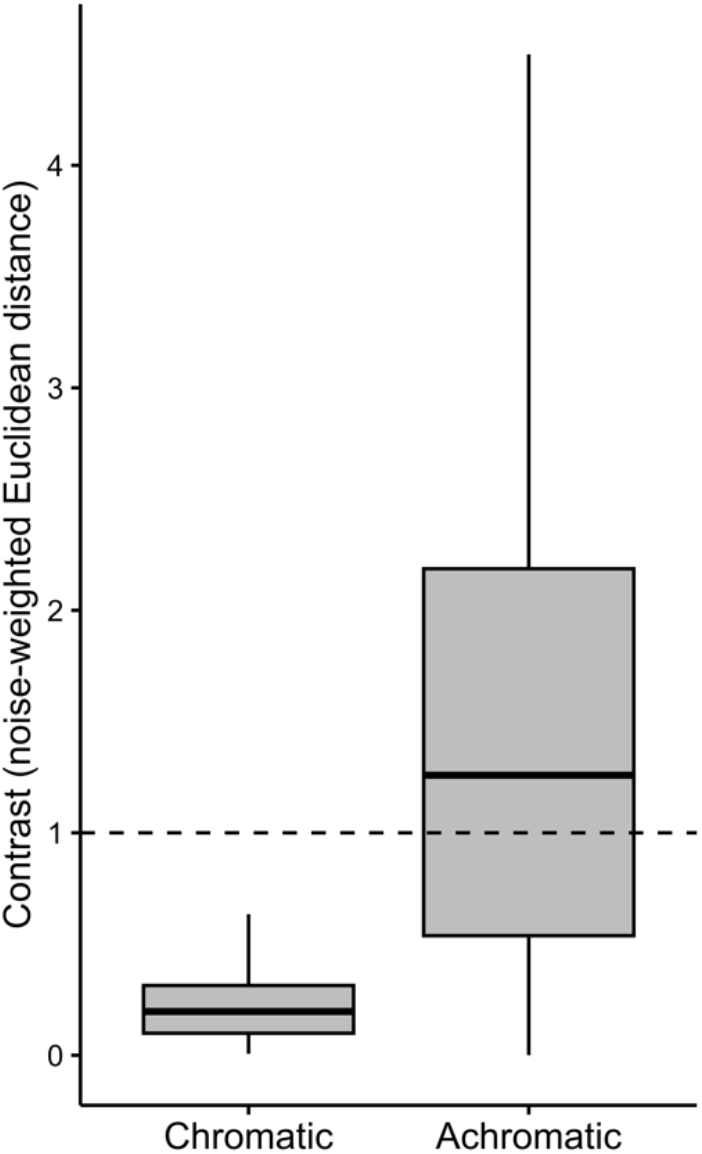
Pairwise chromatic and achromatic contrast between males and females. The dashed line indicates the perceptual threshold above which colors can be distinguished by the avian receiver.

We found a significant negative correlation between male and female chromatic PC1 scores within mated pairs (*r* = -0.60, 95% CI [-0.87–-0.42], t_10_ = -2.38, *P* = 0.04), but there was no significant correlation between paired male and female achromatic PC2 scores (*r* = 0.03, 95% CI [-0.56–0.59], t_10_ = -0.09, *P* = 0.93; Fig. 5). Neither chromatic PC1 (*r* = 0.01, 95% CI [-0.24– 0.25], t_61_ = 0.05, *P* = 0.96) nor achromatic PC2 (*r* = 0.03, 95% CI [-0.22–0.28], t_61_ = 0.24, *P* = 0.81) were correlated with body mass.

**Figure 5:**
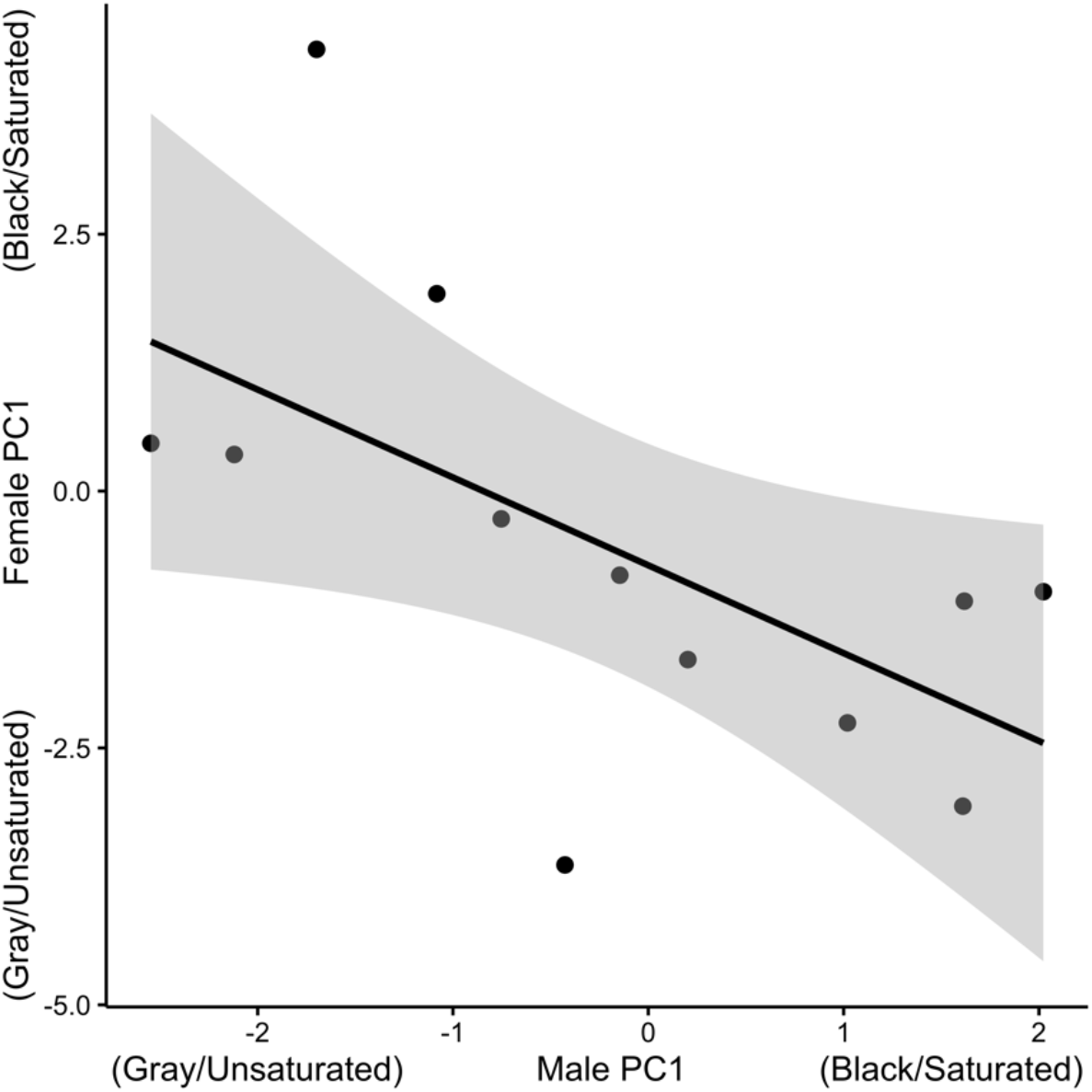
The relationship between male and female chromatic PC1 scores within mated pairs. The solid line represents a linear model fit (shaded area = 95% CI).

## DISCUSSION

Previous studies documented slight but significant sexual dimorphism in Black Terns with regard to morphometric traits (Shealer and Cleary 2007; Shealer *et al*. 2014). Our study is the first to report significant differences in feather reflectance in this species, with breast feathers of males generally darker than females (Figs. 1–3). This difference is not always discernible by human observers in the field (DAS pers. obs.), and our quantitative analyses found considerable overlap between the sexes (Figs. 1–3). Nonetheless, avian visual models suggested that this subtle achromatic difference was sufficient to be distinguished by the receiver (Fig. 4). This conclusion was based on a contrast threshold of 1, but some studies suggest that values between 1 and 3 are difficult to discriminate except under ideal conditions (Siddiqi *et al*. 2004; Langmore *et al*. 2011). However, Black Terns occur in open habitats conducive to visual communication, so it is likely that they can distinguish the sexes based on achromatic plumage variation.

More intriguing, however, was the finding that Black Terns tended to mate disassortatively based on hue and saturation, in that more black and saturated birds were paired with more gray and unsaturated birds (Fig. 5). Morphological studies of mutually ornamented birds have often found positive assortative mating (Jawor *et al*. 2003; MacDougall and Montgomerie 2003; Dias and Cardoso 2023). However, to our knowledge, this is the first report of disassortative mating based on plumage variation in any bird species. Because Black Terns are presumed to be both socially and genetically monogamous, and because mate retention over consecutive years appears to be quite low (Shealer *et al*. 2014), there should be strong selective pressure on traits that advertise individual quality for both males and females. However, a disassortative mating pattern suggests that, if mutual mate choice is occurring, each sex prefers a mate that is in some way dissimilar from itself. It then becomes interesting to consider why this pattern might be manifested in Black Terns and what attributes might be correlated with hue and saturation.

We found no evidence that plumage color was associated with body mass in this system, suggesting it might not be useful as an honest indicator of the signaler’s body condition. Importantly, body mass is a highly plastic trait and there are likely other more relevant indices of body condition that we did not measure. Nonetheless, melanin-based signals such as these may not be greatly affected by condition-dependent mechanisms that could enforce signal honesty (Weaver *et al*. 2017). Such mechanisms are certainly less well understood for melanin-based signals compared to, for example, carotenoid signals (Hill *et al*. 2023). Thus, the fact that melanin-based signals may be less likely to be honest signals – combined with the disassortative mating pattern documented here – suggests that plumage color in this system may serve instead to identify a genetically dissimilar mate (sensu Trivers 1972; Zeh and Zeh 1996). Many studies have documented the benefits of choosing a genetically dissimilar mate, including counteracting the threat of inbreeding depression and increasing offspring immunological diversity, although the overall evidence for this phenomenon in birds is mixed (Mays *et al*. 2007). Future work examining, for example, the genetic relatedness or major histocompatibility complex (MHC) genotypes of mated pairs would help elucidate the potential benefits of mating disassortatively.

Together, these results suggest that Black Terns exhibit plumage variation that is both chromatic and achromatic, but that each component is used in a different context. Achromatic variation subtly distinguishes males from females and thus may be used by individuals to identify sex. Chromatic variation seems more relevant to mate choice, as it is the axis along which males and females mated disassortatively. Not only is the totality of this plumage variation difficult for human observers to distinguish, but the fact that subtle chromatic variation exists that is apparently salient to the birds highlights the importance of analyzing avian visual signals within the context of the appropriate sensory system (Burns and Shultz 2012; Laczi *et al*. 2023). This phenomenon of cryptic phenotypic variation among ostensibly monomorphic and invariable species may be more widespread and warrants further research.

## ACKNOWLEDGEMENTS

Comments from anonymous reviewers greatly improved the manuscript. Fieldwork was carried out by DAS under appropriate trapping, banding, and handling permits issued by the U.S. Bird Banding Laboratory (Master Permit #22827) and Wisconsin Department of Natural Resources. DTB was supported by SUNY Oswego. K. Silvernail assisted with labwork. All applicable ethical guidelines for the use of birds in research have been followed, including those presented in the Ornithological Council’s ‘Guidelines to the Use of Wild Birds in Research’ (Fair *et al*. 2010).

## Notes

### Competing Interest Statement

The authors have declared no competing interest.

